# Pattern component modeling: A flexible approach for understanding the representational structure of brain activity patterns

**DOI:** 10.1101/120584

**Authors:** Jörn Diedrichsen, Atsushi Yokoi, Spencer A. Arbuckle

**Affiliations:** Brain and Mind Institute, Western University, Canada; Department of Statistical and Actuarial Sciences, Western University, Canada; Department of Computer Science, Western University, Canada; Graduate School of Frontier Biosciences, Osaka University, Japan

**Keywords:** Multi-voxel pattern analysis, fMRI, Bayesian models, motor representations

## Abstract

Representational models specify how complex patterns of neural activity relate to visual stimuli, motor actions, or abstract thoughts. Here we review pattern component modeling (PCM), a practical Bayesian approach for evaluating such models. Similar to encoding models, PCM evaluates the ability of models to predict novel brain activity patterns. In contrast to encoding models, however, the activity of individual voxels across conditions (activity profiles) are not directly fitted. Rather, PCM integrates over all possible activity profiles and computes the marginal likelihood of the data under the activity profile distribution specified by the representational model. By using an analytical expression for the marginal likelihood, PCM allows the fitting of flexible representational models, in which the relative strength and form of the encoded features can be estimated from the data. We discuss here a number of different forms with which such flexible representational models can be specified, and how models of different complexity can be compared. We then provide a number of practical examples from our recent work in motor control, ranging from fixed models to more complex non-linear models of brain representations. The code for the fitting and cross-validation of representational models is provided in a open-source Matlab toolbox.

## 1. Introduction

The study of brain representations aims to illuminate the relationship between complex patterns of activity occurring in the brain and "things in the world" - be it objects, actions, or abstract concepts. By understanding internal syntax of brain representations, and especially how the structure of representations changes across different brain regions, we ultimately hope to gain insight into the way the brain processes information.

Central to the definition of representation is the concept of decoding (deCharms and Zador, 2000). A feature (i.e. a variable that describes some aspect of the "things in the world") that can be decoded from the ongoing neural activity in a region is said to be represented there. For example, a feature could be the direction of a movement, the orientation and location of a visual stimulus, or the semantic meaning of a word. Of course, if we allow the decoder to be arbitrarily complex, we would use the term representation in the most general sense. For example, using a computer vision algorithm, one may be able to identify objects based on activity in primary visual cortex. However, we may not conclude necessarily that object identity is represented in V1 - at least not explicitly. Therefore, it makes sense to restrict our definition of an explicit representation to features that can be linearly decoded by a single neuron from some population activity (DiCarlo et al., 2012; DiCarlo and Cox, 2007; Kriegeskorte, 2011; Diedrichsen and Kriegeskorte, 2016).

While decoding approaches are very popular in the study of multi-voxel activity patterns (Haxby et al., 2001; Norman et al., 2006; Pereira et al., 2009), they are not the most useful tool when we aim to make inferences about the nature of brain representations. The fact that we can decode feature X well from region A does not imply that the representation in A is well characterized by feature X - there may be many other features that better determine the activity patterns in this region.

Encoding models, on the other hand, characterize how well we can explain the activities in a specific region using a sets of features. The activity profile of each voxel (here shown as columns in the activity data matrix), is modeled as the linear combination of a set of features (Fig. 1a). We will use the term voxels interchangeably with the more general term measurement channel, which could, depending on the measurement modality, refer to a single neuron, an electrode, or sensor. Each voxel has its own set of parameters **(W)** that determine the weight of each feature. This can visualized by plotting the activity profile of each voxel into the space spanned by the experimental conditions (Fig. 1b). Each dot refers to the activity profile of a channel (here a voxel), indicating how strongly the voxel is activated by each condition. Estimating the weights is equivalent to a projection of each of the activity profiles onto the feature vectors. The quality of the model can then be evaluated by determining how well unseen activity data can be predicted. When estimating the weights, encoding models often use some form of regularization, which essentially imposes a prior on the feature weights. This prior is an important component of the model. It determines a predicted distribution of the activity profiles (Diedrichsen and Kriegeskorte, 2016). An encoding model that matches the real distribution of activity profiles best will show the best prediction performance.

**Figure 1:**
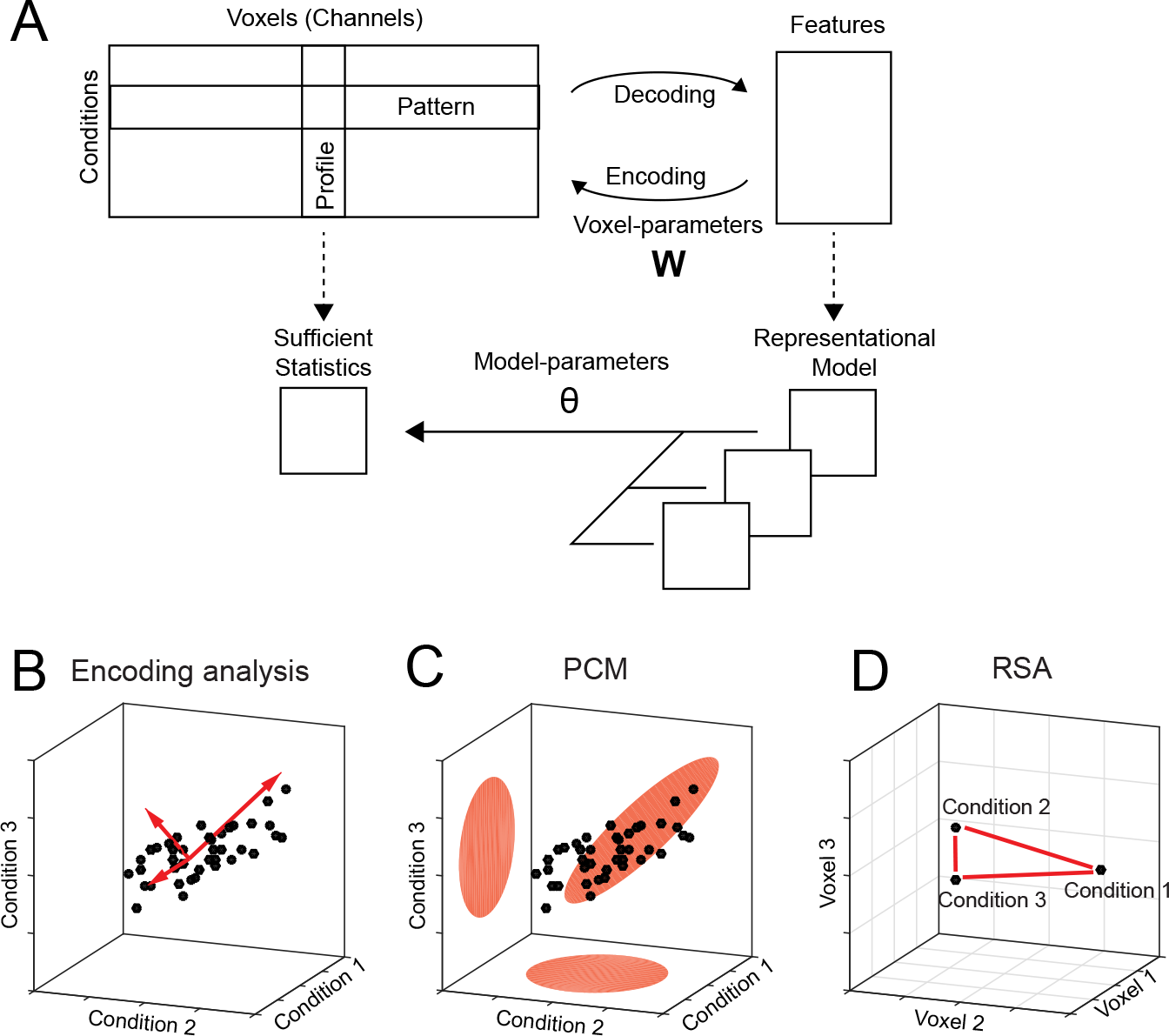
Decoding, encoding and representational models. **(A)** The matrix of activity data consists of rows of activity patterns for each condition or of columns of activity profiles for each voxel (or more generally, measurement channel). The data can be used to decode specific features that describe the experimental conditions (decoding). Alternatively, a set of features can be used to predict the activity data (encoding). Representational models work at the level of a sufficient statistics (the second moment) of the activity profiles. Models are formulated in this space and possibly combined and changed using higher-order model parameters (*θ*). **(B)** The activity profiles of different voxels are plotted as points in the space of the experimental conditions. Features in encoding models are vectors that describe the overall distribution of the activity profiles. **(C)** The distribution can also be directly described using a multivariate normal distribution (PCM).**(D)** Representational similarity analysis (RSA) provides an alternative view by plotting the activity patterns in the space defined by different voxel activities. The distances between activity patterns serves here as the sufficient statistic, which is fully defined by the second moment matrix.

The interpretational problem for encoding models is that for each feature set that predicts the data well, there is an infinite number of other (rotated) features sets that describe the same distribution of activity profiles and, hence, predict the data equally well. The argument may be made that to understand brain representations, we should not think about specific features that are encoded, but rather about the distribution of activity profiles. This can be justified by considering a read-out neuron that receives input from a population of neurons. From the standpoint of this neuron, it does not matter which neuron has which activity profile (as long as it can adjust input weights), and which features were chosen to describe these activity profiles - all that matters is what information can read out from the code. Thus, from this perspective it may be argued that the formulation of specific feature sets and the fitting of feature weights for each voxel are unnecessary distractions.

Therefore, our approach of pattern component modeling (PCM) abstracts from specific activity patterns. This is done by summarizing the data using a suitable summary statistic (Fig. 1a), that describes the shape of the activity profile distribution (Fig. 1c). This critical characteristic of the distribution is the covariance matrix of the activity profile distribution or - more generally - the second moment. The second moment determines how well we can linearly decode any feature from the data. If, for example, activity measured for two experimental conditions is highly correlated in all voxels, then the difference between these two conditions will be very difficult to decode. If however, the activities are uncorrelated, then decoding will be very easy. Thus, the second moment is a central statistical quantity that determines the representational content of the brain activity patterns of an area (Diedrichsen and Kriegeskorte, 2016).

Similarly, a representational model is formulated in PCM not by its specific feature set, but by its predicted second moment matrix. If two feature sets have the same second moment matrix, then the two models are equivalent. Thus, PCM makes hidden equivalences between encoding models explicit. To evaluate models, PCM simply compares the likelihood of the data under the distribution predicted by the model. To do so, we rely on an generative model of brain activity data, which fully specifies the distribution and relationship between the random variables. Specifically, true activity profiles are assumed to have a multivariate Gaussian distribution and the noise is also assumed to be Gaussian, with known covariance structure. Having a fully-specified generative model allows us to calculate the likelihood of data under the model, averaged over all possible values of the feature weights. This results in the so-called model evidence, which can be used to compare different models directly, even if they have different numbers of features. In summarizing the data using a sufficient statistic, PCM is closely linked to representation similarity analysis (RSA), which characterizes the second moment of the activity profiles in terms of the distances between activity patterns (Fig. 1d, also see Diedrichsen and Kriegeskorte 2016).

By removing the requirement to fit and cross-validate individual voxel weights, PCM enables the user to concentrate on a different kind of free parameter, namely model parameters that determine the shape of the distribution of activity profiles. From the perspective of encoding models, these would be hyper-parameters that change the form of the feature or regression matrix. For example, we can fit the distribution of activity profiles using a weighted combination of 3 different feature sets (Fig. 1a). Such component models (see section 2.2.2) are extremely useful if we hypothesize that a region cares about different groups of features (i.e. colour, size, orientation), but we do not know how strongly each feature is represented. In encoding models, this would be equivalent to providing a separate scaling factor to different parts of the feature matrix. Most encoding models, however, use a single model feature matrix, making them equivalent to a fixed PCM model.

In this paper we will discuss the use of PCM to estimate and compare flexible representational models. In section 2, we will present the fundamentals of the generative approach taken in PCM and outline different ways in which flexible representational models with free parameters can be specified. We will then discuss methods for model fitting and for model evaluation. In section 3, we provide three illustrative examples from our work on finger representations in primary sensory and motor cortices, highlighting different representational models and ways of providing inferences on them. The methods presented in this paper are available as a open-source Matlab toolbox (Diedrichsen et al., 2016a).

## 2. Methods

### 2.1. Generative Model

Central to PCM is a generative model of the measured brain activity data **Y**, a matrix of N x P activity measurements, referring to N time points (or trials) and P voxels. The data can refer to the minimally preprocessed raw activity data, or to already deconvolved activity estimates, such as those obtained as beta weights from a first-level time series model. **U** is the matrix of true activity patterns (a number of conditions x number of voxels matrix) and **Z** the design matrix. Also influencing the data are effects of no interest **B** and noise:

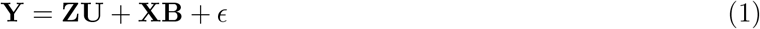

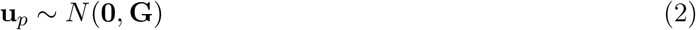

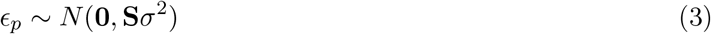

There are a five assumptions in this generative model. First, the activity profiles (u_*p*_, columns of **U**) are considered to be a random variable drawn from a normal distribution. Representational models therefore do not specify the exact activity profiles of specific voxels, but simply the characteristics of the distribution from which they originate. Said differently, PCM is not interested in which voxel has which activity profiles - it ignores their spatial arrangement. This makes sense considering that activity patterns can vary widely across different participants (Ejaz et al., 2015) and do not directly impact what can be decoded from a region. For this, only the distribution of these activity profiles in this region is considered.

The second assumption is that the mean of the activity profiles (across voxels) is the same across conditions, and that it is modeled using the effects of no interests. Therefore, we most often model in **X** the mean of each voxel across conditions. While one could also artificially remove the mean of each condition across voxels (Walther et al., 2016), this approach would remove differences that, from the persepctive of decoding and representation, are highly meaningful (Diedrichsen and Kriegeskorte, 2016).

The third assumption is that the activity profiles come from a multivariate Gaussian distribution. This is likely the most controversial assumption, but it is motivated by a few reasons: First, for fMRI data the multi-variate Gaussian is often a relatively appropriate description, especially if the mean of each voxel across conditions has been removed by the model. Secondly, the definition causes us to focus on the mean and covariance matrix, **G**, as sufficient statistics, as these completely determine the Gaussian. Thus, even if the true distribution of the activity profiles is better described by a non-Gaussian distribution, the focus on the second moment is sensible as it characterizes the linear decodability of any feature of the stimuli.

Fourthly, the model assumes that different voxels are independent from each other. If we used raw data, this assumption would be clear violated, given the strong spatial correlation of noise processes in fMRI data. To reduce these dependencies we typically uses spatially pre-whitened data, which is divided by a estimate of the spatial covariance matrix (Walther et al., 2016; Diedrichsen and Kriegeskorte, 2016). One complication here is that spatial pre-whitening usually does not remove spatial dependencies completely, given the estimation error in the spatial covariance matrix (Diedrichsen et al., 2016b).

Finally, we assume that the noise of each voxel is Gaussian with a temporal covariance that is known up to a constant term σ^2^. Given the many additive influences of various noise sources on fMRI signals, Gaussianity of the noise is, by the central limit theorem, most likely a very reasonable assumption, which is commonly made in the fMRI literature. The original formulation of PCM used a model which assumed that the noise is also temporally independent and identically distributed (i.i.d.) across different trials, i.e. **S = I**. However, as pointed out recently (Cai et al., 2016), this assumption is often violated in non-random experimental designs with strong biasing consequences for estimates of the covariance matrix. If this is violated, we can either assume that we have a valid estimate of the true covariance structure of the noise (*S*), or we can model different parts of the noise structure (see section 2.3).

### 2.2. Representational Model Types

Given our definition (in section 2.1), a representational model is fully specified by the second-moment matrix of the activity profiles **(G).** Many models, including encoding models with a fixed feature set, predict a specific structure of this matrix (fixed models). In other situations we may wish to estimate the most likely structure of **G** from the data without constraints (free models). The most interesting cases, however, are models that impose some constraints on the possible structure of **G**, with the exact form depending on some additional model parameters *θ*.

#### 2.2.1. Fixed models

In fixed models, the second moment matrix **G** is exactly predicted by the model. The simplest and most common example is the Null model, which states that **G = 0.** This is equivalent to assuming that there is no difference between the activity patterns measured under any of the conditions. The Null-model is useful if we want to test whether there are any differences between experimental conditions.

Fixed models also occur when the representational structure can be predicted from some independent data. An example for this is shown in section 3.1, where we predict the structure of finger representations directly from the correlational structure of finger movements in every-day life (Ejaz et al., 2015). Importantly, fixed models only predict the the second moment matrix up to a proportional constant. The width of the distribution will vary with the overall signal-to-noise-level (assuming we use pre-whitened data). Thus, when evaluating fixed models we allow the predicted second moment matrix to be scaled by an arbitrary positive constant.

#### 2.2.2. Component models

A more flexible model is to express the second moment matrix as a linear combination of different components. For example, the representational structure of activity patterns in the human object recognition system in inferior temporal cortex can be compared to the response of a convolutional neural network that is shown the same stimuli (Khaligh-Razavi and Kriegeskorte, 2014). Each layer of the network predicts a specific structure of the second moment matrix and therefore constitutes a fixed model. However, the real representational structure seems to be best described by a mixture of multiple layers. In this case, the overall predicted second moment matrix is a linear sum of the weighted components matrices:

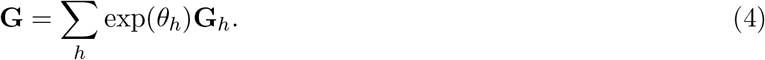

The weights for each component need to be positive - allowing negative weights would not guarantee that the overall second moment matrix would be positive definite. Therefore we use the exponential of the weighing parameter here, such that we can use unconstrained optimization to estimate the parameters.

#### 2.2.3. Feature models

A representational model can be also formulated in terms of the features that are thought to be encoded in the voxels. Features are hypothetical tuning functions, i.e. models of what activation profiles of single neurons could look like. Examples of features would be Gabor elements for lower-level vision models (Kay et al., 2008), elements with cosine tuning functions for different movement directions for models of motor areas (Eisenberg et al., 2010), and semantic features for association areas (Huth et al., 2016). The actual activity profiles of each voxel are a weighted combination of the feature matrix u_*p*_ = Mw_*p*_. The predicted second moment matrix of the activity profiles is then G = MM^*T*^, assuming that all features are equally strongly and independently encoded, i.e. 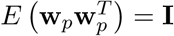. A feature model can now be flexibly parametrized by expressing the feature matrix as a weighted sum of linear components.

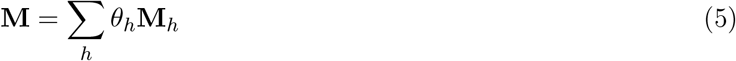

Each parameter *θ_h_* determines how strong the corresponding set of features is represented across the population of voxels. Note that this parameter is different from the actual feature weights **W.** Under this model, the second moment matrix becomes

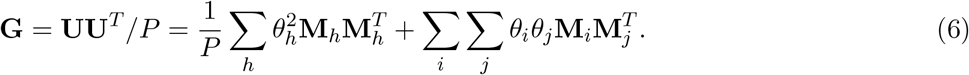

From the last expression we can see that, if features that belong to different components are independent of each other, i.e. M_*i*_M_*j*_ = 0, then a feature model is equivalent to a component model with 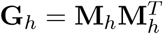. The only technical difference is that we use the square of the parameter *θ_h_*, rather than its exponential, to enforce non-negativity. Thus, component models assume that the different features underlying each component are encoded independently in the population of voxels - i.e. knowing something about the tuning to feature of component A does not tell you anything about the tuning to a feature of component B. If this cannot be assumed, then the representational model is better formulated as a feature model.

#### 2.2.4. Nonlinear models

The most flexible way of defining a representational model is to express the second moment matrix as a non-linear (matrix valued) function of the parameters, **G** = *F* (*θ*). While often a representational model can be expressed as a component or feature model, sometimes this is not possible. One example is a representational model in which the width of the tuning curve (or the width of the population receptive field) is a free parameter (Dumoulin and Wandell, 2008). Such parameters would influence the features, and hence also the second-moment matrix in a non-linear way. Computationally, such non-linear models are not much more difficult to estimate than component or feature models, assuming that one can analytically derive the matrix derivatives ∂**G**/∂*θ_h_*. We will provide an example of this below (see section 3.3).

#### 2.2.5. Free models

The most flexible representational model is the free model, in which the predicted second moment matrix is unconstrained. Thus, when we estimate this model, we would simply derive the maximum-likelihood estimate of the second-moment matrix. This can be useful for a number of reasons. First, we may want an estimate of the second moment matrix to derive the corrected correlation between different patterns, which is less influenced by noise than the simple correlation estimate (Cai et al., 2016; Diedrichsen et al., 2011). Furthermore, we may want to estimate the likelihood of the data under a free model to obtain a noise ceiling - i.e. an estimate of how well the best model should fit the data (see section 2.8).

In estimating an unconstrained **G**, it is important to ensure that the estimate will still be a positive definite matrix. For this purpose, we express the second moment as the square of an upper-triangular matrix, **G = AA**^*T*^ (Cai et al., 2016; Diedrichsen et al., 2011). The parameters are then simply all the upper-triangular entries of **A**.

### 2.3. Noise Model

The noise is assumed to come from a multivariate normal distribution with covariance matrix **S**σ^2^. What is a reasonable noise structure to assume? First, the data can usually be assumed to be independent across imaging runs. If the data are regression estimates from a first-level model, and if the design of the experiment is balanced, then it is usually also permissible to make the assumption that the noise is independent within each imaging run **S = I**, (Diedrichsen et al., 2011). Usually, however, the regression coefficients from a single imaging run show positive correlations with each other. This is due to the fact that the regression weights measure the activation during a condition as compared to a resting baseline, and the resting baseline is common to all conditions within the run (Diedrichsen et al., 2011). To account for this, one can model the mean activation (across conditions) for each voxel with a separate fixed effect for each run. This effectively accounts for any uniform correlation.

Usually, assuming equal correlations of the activation estimates within a run is only a rough approximation to the real co-varince structure. A better estimate can be obtained by using an estimate derived from the design matrix and the estimated temporal autocorrelation of the raw signal. As pointed out recently (Cai et al., 2016), the particular design can have substantial influence on the estimation of the second moment matrix. This is especially evident in cases where the design is such that the trial sequence is not random, but has an invariant structure (where trials of one condition are often to follow trials of another specific condition). The accuracy of our approximation hinges critically on the quality of our estimate of the temporal auto-covariance structure of the true noise. Note that it has been recently demonstrated that especially for high sampling rates, a simple autoregressive model of the noise is insufficient (Eklund et al., 2012).

The last option is to estimate the covariance structure of the noise from the data itself. This can be achieved by introducing random effects into the generative model equation in section 2.1, which account for the covariance structure across the data. One example used here is to assume that the data are independent within each imaging run, but share an unknown covariance within each run, which is then estimated as a part of the covariance matrix (Diedrichsen et al., 2011). While this approach is similar to just removing the run mean from the data as a fixed effect (see above) it is a good strategy if we actually want to model the difference of each activation pattern against the resting baseline. When treating the mean activation pattern in each run as a random effect, the algorithm finds a compromise between how much of the shared pattern in each run to ascribe to the random run-to-run fluctuations, and how much to ascribe to a stable mean activation.

### 2.4. Likelihood and Optimization

Given our generative model we can derive the likelihood of the data under the model. Importantly, we do not want the likelihood for specific values of the the estimates of the true activity patterns **U.** This is a difference to encoding approaches, in which we would estimate the values of **U** by estimating the feature weights **W.** In PCM, we want to assess how likely the data is under any possible value of **U**, as specified by the prior distribution. Thus we wish to calculate the marginal likelihood

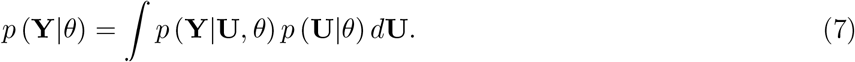

Given that all the involved distributions are normal, this marginal distribution of a single measured activity profile **y**_*i*_ (given the fixed effects estimates **b**_*i*_) can be derived as

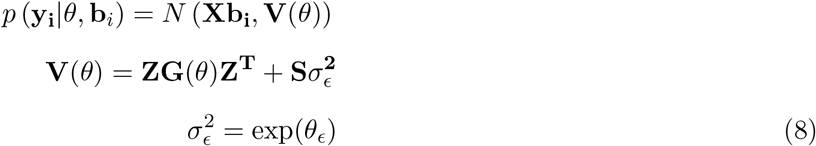

To make our marginal likelihood unconditional on the fixed effects, we need to take into account that we estimate **b**_*i*_ from the data and then use the residuals to calculate the likelihood. This removal can be written as a pre-multiplication with the residual-forming matrix

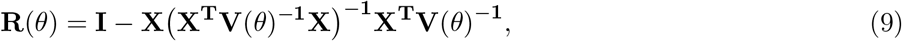
with the residual after removing the fixed effects being **RY**. With this in hand, our final marginal log-likelihood of all data (assuming independence of the voxels) is 
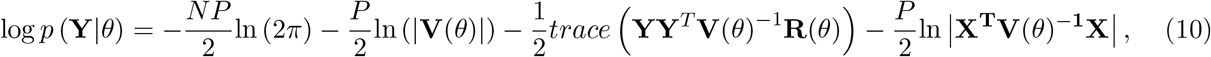
where the last term accounts for the removal of the fixed effects. The interested reader can find a full derivation of the log likelihood and its derivatives in respect to the parameters in the manual accompanying our PCM toolbox. For maximum likelihood estimation, we can use conjugate gradient descent, Newton-Raphson (Lindstrom and Bates, 1988), or Expectation-Maximization (McLachlan and Krishnan, 1997). Implementations of these algorithms are available in the toolbox (Diedrichsen et al., 2016a).

### 2.5. Cross-validated estimate of the second moment matrix

Instead of the full optimization of Eq. 10, we can also obtain simpler estimates of the second moment matrix. The first option is to first estimate the average activity patterns for each condition using linear regression, and then calculate the second moment matrix of these activity patterns.

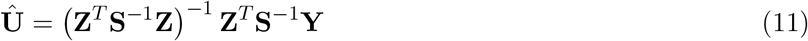

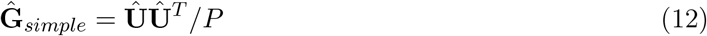

The problem with this naive estimate is that it is positively biased by the measurement noise. In the absence of fixed effects, the expected value of this estimate is

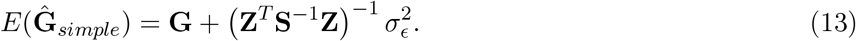

An unbiased estimate of G can be obtained by cross-validation. For each of the independent partitions of the data *m*, we estimate the activity patterns on that run, 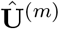 and on all other runs, 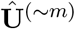 separately. Then, we can calculate the average matrix product over these independent partitions of the data

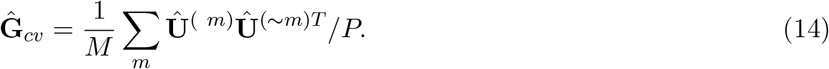

Because in each product we multiply noise terms of independent partitions of the data, we obtain an unbiased estimate of the second moment matrix

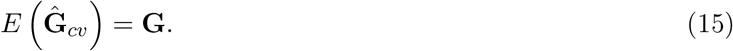

While this estimate of the second moment matrix is unbiased, it is not guaranteed to be positive definite. Thus it is not equivalent to the maximum-likelihood estimate obtained by maximizing Equation 10. For data visualization and as starting values, however, this estimate is extremely useful. Furthermore, distances calculated from this estimate are equivalent to the crossnobis distance estimator, described in detail elsewhere (Diedrichsen and Kriegeskorte, 2016; Walther et al., 2016; Diedrichsen et al., 2016b).

### 2.6. Visualization

After fitting a representational model to the data, it is important to visualize the result. The first obvious visualization is to plot the values of the model parameters *θ*. How important are different feature components to explain the representation in an area?

The second, more comprehensive approach is visualize the second-moment matrix itself. Specifically, we want to compare the fitted second moment matrix to a direct estimate of the second moment matrix from the data. For the latter, we can either use the estimate from a free model, or we can use the cross-validated estimate (Eq. 14). A very powerful way to visualize the second moment matrix is to plot the different conditions into a space spanned by the most important eigenvectors of the second moment matrix (for an example, see Fig. 5 c and d). The distances between different conditions will then show the approximate distances between the patterns. Note that this yields exactly the same visualization as when performing classical multi-dimensional scaling (Borg and Groenen, 2005) on the Euclidean distances between activity patterns (Diedrichsen and Kriegeskorte, 2016).

Finally, we can also visualize the actual estimated activity patterns for each condition across different voxels. This is usually done in encoding approaches, to inspect the spatial distribution of activity profile across the cortical surface. In PCM, the true activation profiles (the random effects) are never explicitly calculated, as they are integrated over when calculating the marginal likelihood (Eq. 7). However, the estimates can be derived as:

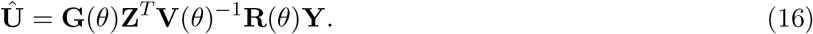

Similarly, for a feature model, the weights for each of the features (columns in the final estimated feature matrix M) can be calculated as^1^

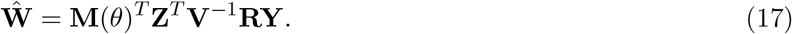

### 2.7. Inference

While visualization is important to further understand the structure of neuronal representations, the main purpose of applying formal models to data is statistical inference. PCM leaves open a number of possibilities here. How to exactly perform inference on a representational model, especially for groups of participants, is still a subject of debate and development. As we will see, the discussion shares a lot of the problems and issues with model inference for other multivariate fMRI models, such as DCM (Penny et al., 2004).

Principally, there are two ways of performing inferences from the fit of a representational model. First, we can perform inference on the parameter estimates. Alternatively we can use the marginal likelihood of the data (Eq. 10) as an approximation of the model evidence to compare which of our candidate models provides the most appropriate description of our data.

#### 2.7.1. Inference based on individual parameter estimates

First we may make inferences based on the parameters of a single fitted model. The parameter may be the weight of a specific component or another metric derived from the second moment matrix. For example, the estimated correlation coefficient between condition 1 and 2 would be 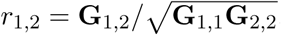. We may want to test whether the correlation between the patterns is larger than zero, or whether a parameter differs between two different subject groups, two different regions, or whether they change with experimental treatments.

The simplest way of testing parameters would be to use the point estimates from the model fit from each subject and apply frequentist statistics to test different hypotheses, for example using a t- or F-test. Alternatively, one can obtain estimates of the posterior distribution of the parameters using MCMC sampling (Murphy, 2012) or Laplace approximation (Friston et al., 2007). This allows the application of Bayesian inference, such as the report of credibility intervals.

One important limitation to keep in mind is that parameter estimates from PCM are not unbiased in small samples. This is caused because estimates of **G** are constrained to be positive definite. This means that the variance of each feature must be larger or equal to zero. Thus, if we want to determine whether a single activity pattern is different from baseline activity, we cannot simply test our variance estimate (i.e. elements of **G)** against zero - they trivially will always be larger, even if the true variance is zero. Similarly, another important statistic that measures the pattern separability or classifiability of two activity patterns is the Euclidean distance, which can be calculated from the second moment matrix as *d* = **G**_1,1_ + **G**_2,2_ **−** 2**G**_1,2_. Again, given that our estimate of **G** is positive definite, any distance estimate is constrained to be positive. To determine whether two activity patterns are reliabily different, we cannot simply test these distances against zero, as the test will be trivially larger than zero. A better solution for inferences from individual parameter estimates is therefore to use a cross-validated estimate of the second moment matrix (Eq. 14) and the associated distances (Walther et al., 2016; Diedrichsen et al., 2016b). In this case the expected value of the distances will be zero, if the true value is zero. As a consequence, variance and distance estimates can become negative. These techniques, however, take us out of the domain of PCM and into the domain of representational similarity analysis (Kriegeskorte et al., 2008; Diedrichsen and Kriegeskorte, 2016).

#### 2.7.2. Inference using model evidence

As an alternative to parameter-based inference, we can fit multiple different models and then compare them according to their model evidence the likelihood of the data given the models (integrated over all parameters). In encoding models, the weights **W** are directly fitted to the data, and hence it is important to use cross-validation to compare models with different numbers of features. The marginal likelihood (Eq. 10) already integrates all over all likely values of **U**, and hence **W**, thereby removing the bulk of free parameters. Thus, in practice the marginal likelihood will be already close to the true model evidence.

Our marginal likelihood (Eq. 10), however, still depends on the free parameters *θ*. So, when comparing models, we need to still account for the risk of overfitting the model to the data. For fixed models, there are only two free parameters: one relating to the strength of the noise and one relating to the strength of the signal. This compares very favorably to the vast number of free parameters one would have in an encoding model, which is the size of **W**, the number of features x number of voxels. However, even the fewer model parameters still need to be accounted for. We propose here four ways of doing so.

The first option is to use empirical Bayes or Type-II maximal likelihood. This means that we simply replace the unknown parameters with the point estimates that maximize the marginal likelihood. This is in general a feasible strategy if the number of free parameters is low and all models have the same numbers of free parameters, which is for example case when we are comparing different fixed models. The two free parameters here determine the signal-to-noise ratio. For models with different numbers of parameters we can penalize the likelihood by 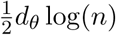, yielding the Bayes information criterion (BIC) as the approximation to model evidence.

As an alternative option, we can use cross-validation within the individual (Fig. 2a) to prevent overfitting for more complex flexible models, as is also currently common practice for encoding models (Naselaris et al., 2011). Taking one imaging run of the data as test set, we can fit the parameters to data from the remaining runs. We then evaluate the likelihood of the left-out run under the distribution specified by the estimated parameters. By using each imaging run as a test set in turn, and adding the log-likelihoods (assuming independence across runs), we thus can obtain a approximation to the model evidence. Note, however, that for a single (fixed) encoding model, cross-validation is not necessary under PCM, as the activation parameters for each voxel **(W** or **U)** are integrated out in the likelihood. Therefore, it can be handled with the first option we described above.

**Figure 2:**
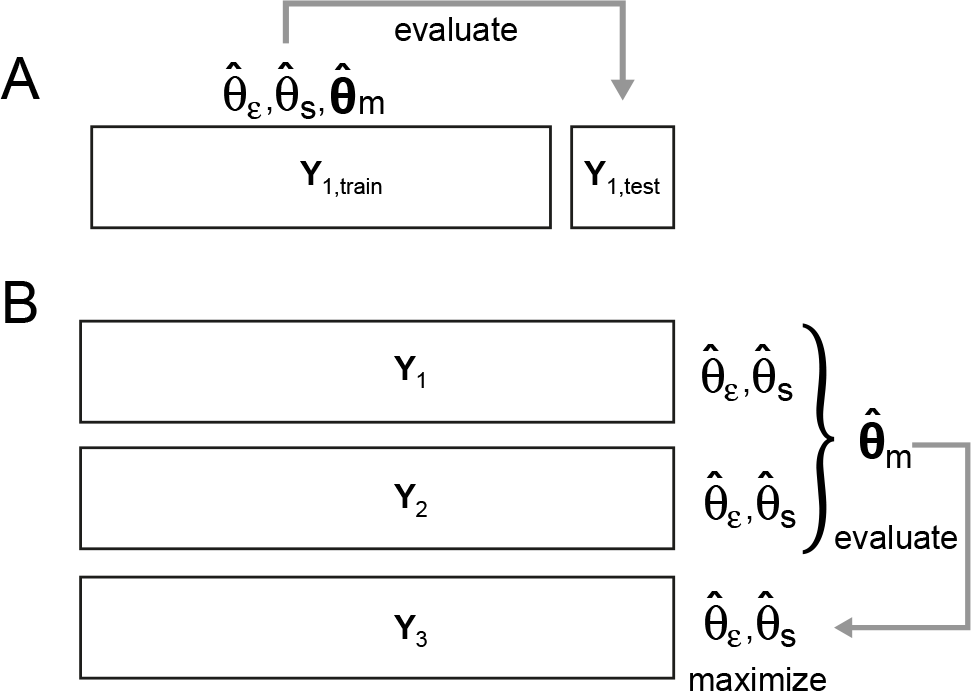
Cross-validation for testing representational models. (**A**) Within-subject cross-validation. Both the model parameters (*θ_m_*), as well as the noise (*θ_ε_*), and the signal parameter (*θ_s_*) are fitted on the training data set. The likelhood of the test data (**Y**_1,*test*_) is then evaluated under those parameters. (**B**) Group crossvalidation, here shown with the data from three subjects (**Y**_1–3_). The model parameters are commonly fit to all but one subject, allowing each subject a separate noise and signal parameter. The likelihood of the data from the left-out subject is then evaluated under the group model parameters, but optimizing the individual’s noise and signal parameter.

For the third option, if we want to test the the hypothesis that the representational structure in the same region is similar across subjects, we can perform cross-validation across participants (Fig. 2b). We can estimate the parameters that determine the representational structure using the data from all but one participants and then evaluate the likelihood of data from the left-out subject under this distribution. When performing cross-validation within individuals, a flexible model can fit the representational structure of individual subjects in different ways, making the results hard to interpret. When using the group cross-validation strategy, the model can only fit a structure that is common across participants. Different from encoding models, representational models can be generalized across participants, as we do not fit the actual activity patterns, but rather the representational structure. In a sense, this method is performing “hyper alignment” (Guntupalli et al., 2016) without explicitly calculating the exact mapping into voxel space (Eq. 17). When using this approach, we still allow each participant to have it’s own signal and noise parameters, because the signal-to-noise ratio is idiosyncratic to each participant’s data. When evaluating the likelihood of left-out data under the estimated model parameters, we therefore plug in the ML-estimates for these two parameters for each subjects.

Finally, a last option is to implement a full Bayesian approach and to impose priors on all parameters, and then use a Laplace approximation to estimate the model evidence (Kass and Raftery, 1995; Friston et al., 2007). While it certainly can be argued that this is the most elegant approach, we find that cross-validation at the level of model parameters provides us with a practical, straightforward, and transparent way of achieving a good approximation.

Each of the inference strategies supplies us with an estimate of the model evidence. To compare models, we then calculate the log Bayes factor, which is the difference between the log model evidences.

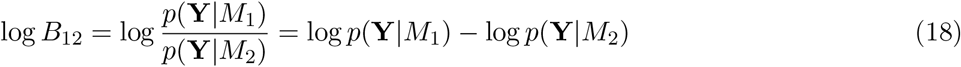

Log Bayes factors of over 1 are usually considered positive evidence for one model over the other (Kass and Raftery, 1995).

#### 2.7.3. Performing group inference on representational models

How to perform group inference in the context of Bayesian model comparison is an topic of ongoing debate in the context of neuroimaging. A simple approach is to assume that the data of each subject is independent (a very reasonable assumption) and that the true model is the same for each subject (a maybe less reasonable assumption). This motivates the use of log Group Bayes Factors (GBF), which is simple sum of all individual log Bayes factor across all subjects n

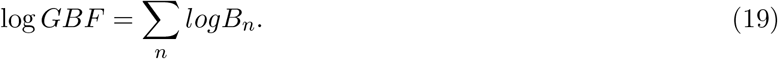

Performing inference on the GBF is basically equivalent to a fixed-effects analysis in neuroimaging, in which we combine all time series across subjects into a single data set, assuming they all were generated by the same underlying model. A large GBF therefore could be potentially driven by one or few outliers. We believe that the GBF therefore does not provide a desirable way of inferring on representational models - even though it has been widely used in the comparison of DCM models (Friston et al., 2003).

At least the distribution of individual log Bayes factors should be reported for each model. When evaluating model evidences against a Bayesian criterion, it can be useful to use the average log Bayes factor, rather than the sum. This stricter criterion is independent of sample size, and therefore provides a useful estimate or effect size. It expresses how much the favored model is expected to perform better on a new, unseen subject. We can also use the individual log Bayes factors as independent observations that are then submitted to a frequentist test, using either a t-, F-, or nonparametric test. This provides a simple, practical approach that we will use in our examples here. Note, however, that in the context of group-crossvalidation, the log-Bayes factors across participants are not strictly independent.

Finally it is also possible to build a full Bayesian model on the group level, assuming that the winning model is different for each subject and comes from a multinomial distribution with unknown parameters (Stephan et al., 2009).

### 2.8. Noise ceilings

Showing that a model provides a better explanation of the data as compared to a simpler Null-model is an important step. Equally important, however, is to determine how much of the data the model does not explain. Noise ceilings (Nili et al., 2014) provide us with an estimate of how much systematic structure (either within or across participants) is present in the data, and what proportion is truly random. In the context of PCM, this can be achieved by fitting a fully flexible model, i.e. a free model in which the second moment matrix can take any form. The non-crossvalidated fit of this model provides an absolute upper bound - no simpler model will achieve a higher average likelihood. As this estimate is clearly inflated (as it does not account for the parameter fit) we can also evaluate the free model using cross-validation. Importantly, we need to employ the same cross-validation strategy (within /between subjects) as used with the models of interest. If the free model performs better than our model of interest even when cross-validated, then we know that there are definitely aspects of the representational structure that the model did not capture. If the free model performs worse, it is overfitting the data, and our currently best model provides a more concise description of the data. In this sense, the performance of the free model in the cross-validated setting provides a “lower bound” to the noise ceiling. It still may be the case that there is a better model that will beat the currently best model, but at least the current model already provides an adequate description of the data. Because they are so useful, noise ceilings should become a standard reporting requirement when fitting representational models to fMRI data, as they are in other fields of neuroscientific inquiry already. The Null-model and the upper noise ceiling also allow us to normalize the log model evidence to be between 0 (Null-model) and 1 (noise ceiling), effectively obtaining a Pseudo-*R*^2^.

## 3. Examples

In this section we provide examples of how to build and evaluate representational models from our recent studies on movement representations. We did not always use PCM in the original papers - partly because some of the methods were not fully developed at this point - partly because other strategies of getting similar results (i.e. using RSA with cross-validated distances) were at this point more common and easier to communicate. We show here how to perform these analyses using PCM in a concise manner - often allowing for more powerful inferences.

### 3.1. Representational structure of finger movements

In a recent paper (Ejaz et al., 2015), we studied the activity patterns associated with single finger movement in primary sensory and motor cortices. The actual patterns of neural activity were quite very variable across different participants. However, when we studied the second moment matrix of the activity profile distribution (Fig. 3a), we found that they were highly correlated across different individuals. As can be seen from a multi-dimensional scaling of the activity patterns (Fig. 3b), the thumb (d1) showed the most unique pattern, while the patterns of other fingers (d2-d5) were arranged according to their neighborhood relationship. Interestingly, the pattern for the little finger (d5) was already more similar to the thumb pattern than the middle or ring finger - thus the structure was not well explained by a simple somatotopic ordering of the fingers on the cortical sheet.

**Figure 3:**
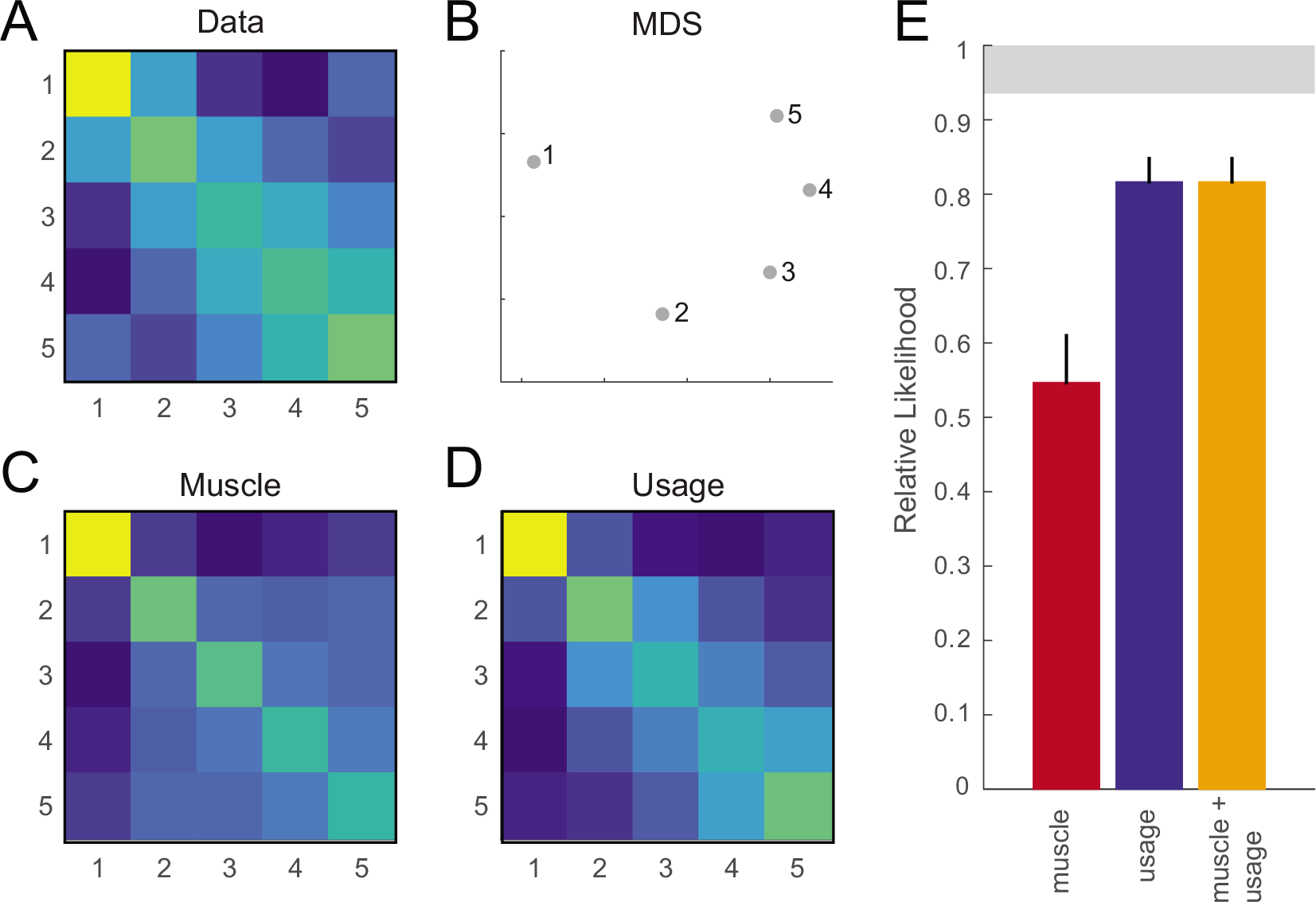
Comparison of representational models of the structure of finger representations using PCM. (**A**) A cross-validated estimate of the second moment of the activity profile distribution. The rows and columns of the matrix refer to digit 1-5. (**B**) Classical multi-dimensional scaling of the representational structure. Shown are the finger patterns projected on the eigenvectors of the second moment matrix (**C**) Predicted second-moment matrix under the muscle model. (**D**) Predicted second-moment matrix under the natural statistics of hand movements. (**E**) Pseudo-R2 based on the log-Bayes factor for the models against a null model. The gray bar indicates the area between the lower and upper noise ceiling.

What can explain this highly invariant representation? We tested a number of hypotheses. First, we considered the possibility that two cortical activity patterns could also be more similar, because the two movements involved similar sets of muscles. We therefore measured the muscle activity involved in making the single finger movements and predicted the second-moment matrix of the patterns directly from the correlation matrix of the measured muscle patterns (Fig. 3c). Alternatively, we measured the movements of the fingers during normal every-day activities (Ingram et al., 2008), hypothesizing that two fingers that commonly move together would also exhibit similar activity pattern. Thus, we predicted the second-moment matrix of the patterns directly from the correlation matrix of the natural statistics of finger movements (Fig. 3d).

In the original paper (Ejaz et al., 2015), we used RSA to compare the models. However, we have recently shown that the same model comparison can be performed in a more powerful fashion through PCM (Diedrichsen and Kriegeskorte, 2016). The two models give us a classic example of “fixed” models, in which we fully predict the distribution of activity profiles (up to a signal scaling constant) using external data. Per subject, we therefore only estimated the signal and noise parameter to optimize the likelihood (Eq. 10). Because both models had only those two free parameters, we used Type-II maximum likelihood (empirical Bayes) as an estimate of model evidence.

To scale the model evidence we also fit a null model. In this case the null model predicted that the second moment matrix would be identity - that is, all finger patterns are equally far apart from each other. We then expressed each subject’s model evidence relative to the null model by taking the difference between log model evidences (see section 2.7.2). Note that we also could have used the null model that there are no differences between activity patterns. This would have given us much larger log-Bayes factors against the null model, but would have left the differences between models unchanged.

We also calculated a noise ceiling by fitting a free model of the the second moment matrix either to all the subjects (upper bound) or to all participants, excluding the one that we evaluated the fit on (lower bound, see section 2.8).

We can then display the model evidence in the form of a “Pseudo-R2”, with zero referring to the null model and 1 to the upper noise ceiling. As can be seen from Figure 3e, the usage model was consistently associated with a larger evidence than the muscle model. This was the case in all of the seven participants. The average log-Bayes factor for a single individual was 81.56 (SE=24.801), suggesting overwhelming evidence for the usage over the muscle model. The usage model did not, however, reach the lower noise ceiling, indicating that there is still systematic structure in the representations that is unexplained by the model.

We then explored whether a combination of the two models would explain the data better than the hand usage model alone. Thus, we built a component model, where the two component matrices (Eq. 4) corresponded to predicted second moment matrices from the muscle and usage model, respectively. Thus, we are exploring the idea that some voxels in our population may be better described as tuned to muscle activations independent of natural usage, whereas others reflect the natural statistics, with the exact proportions of these populations unknown. While we here only combine 2 components, in ongoing work we are using many more components to describe the representation of movement sequences.

Adding more possible components will of course lead to a better fit to the data. Thus, to evaluate the model evidence, we are using here crossvalidation across subjects (Fig 2b), estimating the mixture proportions on data from all but one subject and then evaluating the fit on the left-out subject (only allowing free scaling and noise parameters, as for the other models). In this particular case, the combination model did not perform better than the usage model alone. The weight of the muscle model was usually very low, and the log Bayes factor between the two models was ever so slightly in favour of the simpler usage model, indicating that the combination model overfit the data.

### 3.2. Correlation between ipsilateral and contralateral finger presentations

The common application of PCM (and the one that it was originally developed for) is to estimate the true correspondence between two activity patterns. For example, both movements of the fingers of the contralateral and ipsilateral hand evoke activity patterns in primary motor cortex that allow one to decode which finger moved (Diedrichsen et al., 2013). The patterns evoked by ipsilateral movement are weaker as compared to the patterns elicited by contralateral movement, but they appear to match on a finger-by-finger basis.

To estimate the extent of this correspondence, we can estimate the covariance matrix between the pattern for the contra (c) and ipsilateral (i) finger and then calculate the correlation between a matching finger pair as

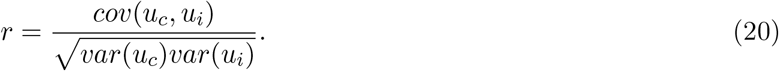

For such a correlation to be informative, we first need to subtract the mean activity patterns for all fingers of each hand, to ensure that the correlations are not simply driven by a correlation between the overall activity patterns. Using this simple estimate of the covariance (Eq. 11) between finger-specific activity patterns, we obtain an average correlation between finger pairs of *r =* 0.28(+0.02), which is significantly larger than zero. This measure however, does not give us any information about the true degree of correspondence, as the simple covariance estimate is biased by measurement noise (see Eq. 13). This fact causes our correlation estimates to be much smaller than 1, even if the true correlation was 1.

Using PCM, we can model a finger-by-finger correlation using a feature model with three parameters (Fig. 4b). The patterns of the contra-lateral hand are modeled to consists of the finger specific patterns **w**_1–5_, each weighted by the parameter *θ*_*a*_. Note, that while each finger is associated with it’s own finger pattern, all fingers share the same weighting parameter. This means that under this model all finger patterns of the contralateral hand have the same variance 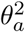, as can be seen from the predicted second moment matrix (Fig. 4c). The patterns for the ipsilateral fingers are modeled as the sum of the contralateral pattern, weighted by *θ_b_*, and an independent ipsilateral finger-specific pattern (**W**_6–10_), weighted by *θ_c_*. Accordingly, the variance of the ipsilateral patterns will be 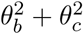, and the covariance *θ_a_θ_b_*. If the ipsilateral patterns are only a scaled version of the contralateral patterns (*θ_c_* = 0), the correlation will be 1. Indeed, when we estimated the model parameters on the real data, we obtain correlations estimates of 1 in 11 of the 12 cases (Fig. 4d).

**Figure 4:**
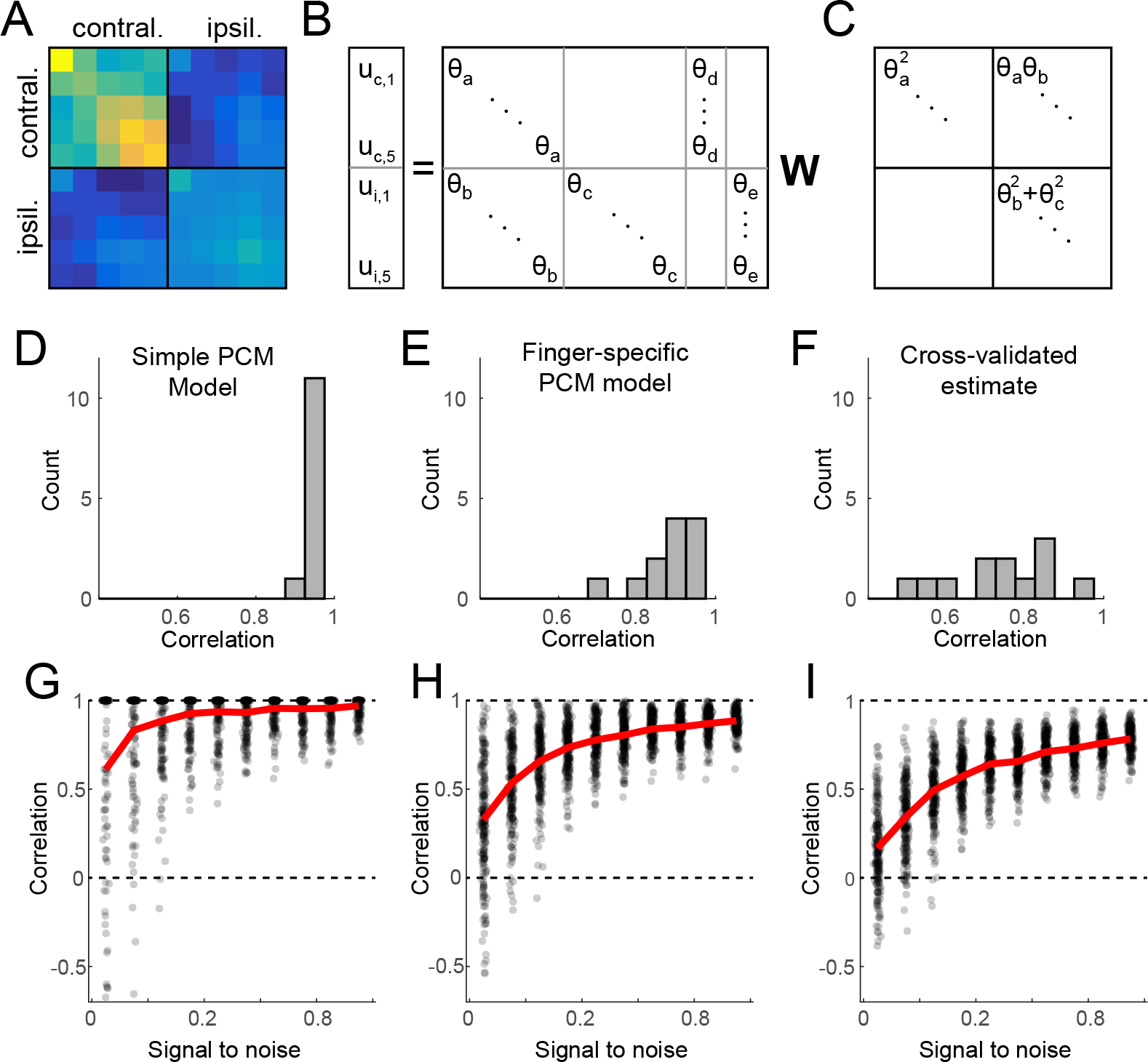
Estimating the true correlation between activity patterns. **(A)** Crossvalidated estimate of the covariance matrix of fingers of the contra- and ipsilateral hands. The upper right 5x5 panel indicates the co-variances of finger patterns across hands. The fact that the diagonal of this part of the matrix is higher than the off-diagonal, indicates a positive correlation of matching finger pairs. **(B)** Feature model, expressing the patterns of the contra-lateral fingers (*u*_*c*,1_ − *u*_*c*,5_) and the patterns of the ipsilateral fingers (*u*_*i*,1_ − *u*_*i*,5_) as a combination of the shared finger pattern (weighted by *θ_a_* and *θ_b_*, respectively) and one that is idiosyncratic to the ipsilatal hand, weighted by *0*_*c*_. Empty parts of the matrix are zero. To complete the model, we also added a hand-specific pattern for the contra and ipsilateral hand, weighted by *θ_d_* and *θ_e_* respectively **(C)** Under the assumption that the rows of **W** have variance of 1 and are uncorrelated, the predicted second moment matrix of the activity profiles can be derived (expressed here ignoring the common hand patterns). **(D)** Distribution of the estimated correlation coefficients for data from 12 hemispheres, using the feature model. **(E)** Distribution of the estimated correlation coefficients using a feature model the allows for finger-specific variances. **(F)** Distribution of estimated correlation coefficients using the cross-validated estimate of the second moment matrix. **(G)** Simulation assuming that the contra- and ipsilateral patterns are perfectly correlated (r=1), but the signal variance across hands and fingers are uneven. Correlation estimates approach 1 as signal-to-noise ratio (x-axis) increases. **(H)** The same simulation for finger-specific feature model and **(I** for cross-validated estimates. The dots reflect the estimated correlation from single simulations. Red lines indicate the average.

The model restricts the finger-specific variances to be the same across all finger, something that is definitely not the case when inspecting the estimated second moment matrix (Fig. 4a). In the original paper, we therefore allowed each finger to have its own three parameters to describe variances and covariance of the contra- and ipsilateral patterns. From these, we calculated the average variances and covariances across fingers, before calculating the correlation. This approach led to smaller correlation estimates (Fig. 4e). To determine which of these approaches provides more realistic estimates, we simulated data at different signal-to-noise levels using unequal variances across hands (1 and 0.2), unequal variances for different fingers (1, 0.5, 0.1), and a correlation of 1. When applying the two models, we can see that the simpler model that forces all fingers to have the same variance (Fig. 4g) shows less bias than the model that allowed for finger-specific variances (Fig. 4h). Thus, these analysis show that a highly structured model designed to directly recover the parameter of interest can be superior to an approach in which the covariance matrix is modeled flexibly and then averaged to retrieve the quantities of interest.

An alternative, possibly simpler, way of accounting for the noise, is to use a cross-validated estimate of the covariance matrix (Eq. 14, Fig. 2b). This estimate of the covariance matrix is unbiased, but is not guaranteed to be positive definite (see section 2.5). When calculating correlations based on such covariance estimates, one can encounter a number of problems. If one of the variance estimates (diagonal entries in the matrix) is negative, the correlation is undefined. Even if the variance estimates are forced to be positive the covariance might exceed the denominator of Eq. 20, leading to correlations larger than 1 or smaller than -1. To make the correlation estimate more stable, we can force the matrix to be semi-positive definite by removing eigenvectors with negative eigenvalues. Using this technique, we obtain smaller estimates of the correlation (Fig. 4f). Simulations indeed show that if the true correlation between finger pairs is 1, the estimates are substantially biased towards smaller values (Fig. 4i).

Thus, cross-validation cannot provide us with good estimates of the true correlation between two patterns. Even though PCM estimates are still biased, the estimates are more accurate than those using a other approaches. Based on our results, we can therefore draw strong conclusions about the bilateral representations of finger movements: the finger-specific activity patterns that occur during ipsilateral finger movements are weaker echoes of the patterns evoked by movement of the corresponding contral-ateral finger. There is no evidence in the data of any pattern component that would be unique to the ipsilateral movement, which would be expected if the area performed a specific function in ipsilateral movements that was distinct from its contribution to the contralateral movements.

### 3.3. Changes in finger patterns with movement speed

In an ongoing experiment, we are studying how the representations of single finger movements change when the frequency of finger movement increases. The answer to this question is important to understand how patterns of neural activity related to patterns of fMRI activity - and whether this mapping changes when the overall BOLD activity of the region increases. ECoG recording over human sensory-motor cortices have shown that the cumulative neural activity in the high-frequency band (HFB) increases non-linearly with the tapping frequency (Hermes et al., 2012), mostly related to the fact that neural activity of subsequent presses is suppressed relative to the first press. At the same time, the average BOLD signal in primary motor cortex appears to be roughly linearly related to HFB activity (Siero et al., 2013). Would this also be true on the pattern level? If yes, we would expect that the finger patterns would just scale relative to baseline with increasing tapping speed - but that their relative arrangement would remain stable across tapping speeds.

We can formalize this prediction into a model, in which the relationship between finger patterns in captured in a free model, with 15 parameters that describe the variances and co-variance between the fingers (**A**, Fig. 5a). This representational structure would then be scaled by a separate speed-dependent constant *θ_a–c_* in matrix **M**. To remove parameter redundancy, we fix the scaling factor for the highest speed to 1. Under this model, the predicted second movement matrix would take a form similar to that depicted in Figure 5b. When plotting this prediction in representational space (Fig. 5c and d), one can see that the distances between finger patterns increase with increasing speed, while their overall arrangement remains stable. More precisely, the activity patterns should just increase linearly from rest. Finding such a pattern would provide strong evidence for the linearity of th BOLD signal to neural activity even on the pattern level.

The model is non-linear, as overall feature matrix depends on a product of two variables. The derviates of the second moment matrix can be obtained following the chain and product rules for matrix derivatives.

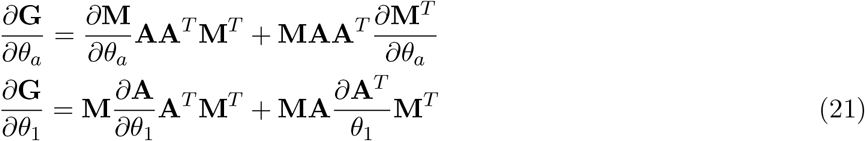

**Figure 5:**
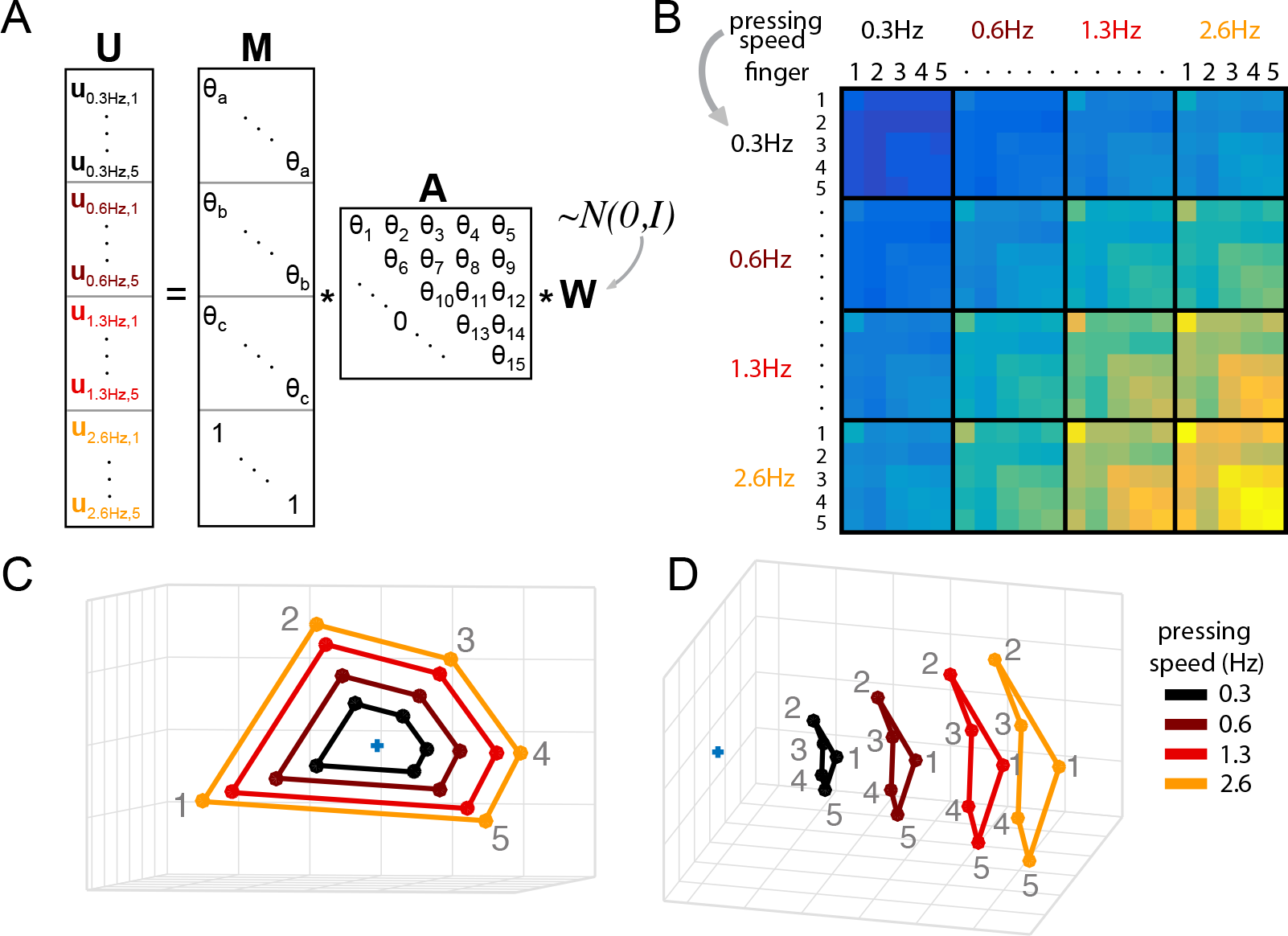
Scaling model of finger activity patterns. (**A**) Model structure. The activity patterns **U** are scaled for each pressing speed by an arbitrary scaling parameter *θ_a–c_* defined in matrix **M**. The off-diagonals are zero (not shown). The covariances between finger patterns (across speeds) are modeled here using a free model with 15 coefficients defined in the upper-triangular matrix **A**. (**B**) Predicted second-moment matrix of the scaling model. The 5x5 block along the diagonal show the covariances of finger patterns elicited at one of four pressing speeds (each speed is denoted by a different color). Off-diagonal panels indicate the covariances between finger patterns measured from different pressing speeds. (**C,D**) Classical multi-dimensional scaling of the predicted representational structure from the predicted covariance matrix. Each finger pattern is scaled by a speed-dependent constant relative to rest. Colors indicate pressing speed, numbers indicate fingers (1-5). The blue crosses indicate resting baseline.

This enables us to fit the model to the data in a computationally efficient manner.

## 4. Discussion

In this paper we have presented a statistical approach for fitting and comparing representational models. PCM is linked very closely to both encoding models and RSA (Diedrichsen and Kriegeskorte, 2016), but the particular approach makes it especially attractive for the fitting and comparison of flexible representational models.

In encoding models, a separate weight of each voxel and feature is explicitly fitted. To evaluate model fit, the prediction accuracy for left-out data is used. In contrast, PCM evaluates the marginal likelihood of the data under the distribution predicted by the model. In this expression, the unknown activity profiles for each voxel are integrated out (Eq. 7). This means that we can evaluate different encoding models, even with different numbers of features, by simply comparing their marginal likelihoods. The step of integrating over the actual activity profiles is justified under the assumption that the exact spatial arrangement of the different activity profiles does not matter for the representational content of a brain area - even if the neurons were differently arranged, a fully connected read-out neuron could extract the same information. Indeed, it can be shown that the spatial arrangement of patterns associated with finger movements is variable in primary motor cortex, whereas the representational structure - and hence likely the function -is remarkably invariant across participants (Ejaz et al., 2015).

PCM uses the second moment as a summary statistics for observed data. Analogously, it also uses the second moment to specify the representational model. It therefore constitute and abstraction from the use of concrete feature sets. Using features to describe a population of activity profiles has a long tradition in Neuroscience. In motor control, the response properties of single neurons in primary motor cortex have often been characterized as being tuned to movement direction (Georgopoulos et al., 1986). However, many studies have made clear that representations in M1 contain a complex mixture of features, including position (Sergio and Kalaska, 1997), direction (Schwartz et al., 1988), force (Sergio et al., 2005), or muscle activity (Kakei et al., 1999), often with non-linear interactions between them (Sergio and Kalaska, 1997). Indeed, it has been questioned whether a description of population activity in terms of semantically definable features is at all sensible (Shenoy et al., 2013).

This is also one of the key lessons from the emerging field of deep learning algorithms as a model of neural processing (Marblestone et al., 2016). Most often, the learned features found in the hidden layers defy simple semantic description. This means that we need tools to formulate and test hypotheses about neural representations that describe the representational space itself, without getting distracted by the relatively superficial issue of which axes should be used to describe the space. Indeed, the representational structure found in different layers of a deep neural networks can be used as a model for representational structures found in the human brain (Khaligh-Razavi and Kriegeskorte, 2014). Even if it turns out that real neural representations are distinctively different from the activation states found in artificial neural networks, deep nets form an ideal testing ground for the methods we employ to understand biological systems. If our methods cannot provide insight into the transformations occurring between the layers of these still relatively simple networks, they most likely will tell us very little about what happens in the brain (Jonas and Kording, 2017).

PCM has a very tight relationship to RSA, in which we also use a summary statistics to compare data and model predictions. In the case of RSA, the summary statistics are the distances between all pairs of neural activity patterns. These distances usually contain the same information as the second moment matrix. Furthermore, RSA has also been used to fit flexible representational models (Khaligh-Razavi et al., 2017). However, unless one takes into account the co-dependences between these distance estimates, PCM provides more powerful inferences as long as its assumptions are approximately satisfied (Diedrichsen and Kriegeskorte, 2016).

Not having to fit and cross-validate the individual voxels feature weights makes it easier to explore a larger space representational models. Note that a simple encoding model (despite all the free feature weight parameters) is a fixed PCM model, in which the only signal and noise parameter needs to be fitted. More complex models, in which the second moment matrix is to some degree flexible, would refer to extension, differential scaling, or nonlinear changes of the model features. Having an analytical expression for the marginal likelihood allows us to directly estimate these parameters. For model comparison, we can use simple cross-validation (rather than the double cross-validation necessary in encoding approaches). It is also possible to avoid cross-validation completely, by using a fully Bayesian approach, in which a normal prior is imposed on the parameters *θ*. Here we chose cross-validation as a practical and robust approach for estimating the model evidence.

Despite these practical advantages, there are a number of drawbacks and limitations to our approach. The reader should be cautioned that, although PCM attempts to separate the contribution of signal and noise to the second moment of the data, the estimate of the true second moment matrix is not unbiased when using small samples. This is due to the fact that the estimate of **G** is constrained to be positive definite. Hence, when the true signal variance is zero, the variance estimates will be zero or positive, causing the mean estimates to be larger than zero. For unbiased results, one would need to use cross-validation (Eq. 14), which sometimes also results in negative variance estimates. Note, however, that although the cross-validated second moment matrix estimate is unbiased, correlation coefficients derived from this estimate are not. In contrast, correlation estimates derived from PCM have a smaller bias. In general, however, the restriction to positive definite estimates prevents us from being able to determine, based on the second moment matrix alone, whether a certain feature is encoded above chance. For this, we would need to compare a representational model with and without that feature encoded (see section 3.1).

Another limitation of PCM arises from the fact that it uses only the second moment of the data as a sufficient statistic. In this, PCM ignores certain aspects of the neural code. First, it neglects the spatial arrangement of the different activity profiles on the cortical sheet. One of the coding principles in the neocortex is that nearby locations also have similar activity profiles (Graziano and Aflalo, 2007). Therefore, the exact distribution of activity profiles across the cortical surface may contain some important insights into information coding. On the other hand, there is evidence that the exact arrangement of activity patterns on the cortical surface may reflect random biological variation that in itself does not speak to the computational function of the region (Ejaz et al., 2015).

Similarly, PCM ignores all higher statistical moments of the activity profile distribution. In primary visual cortex, for example, many voxels are tuned to one specific stimulus locations in the visual field, but very few respond to two separate locations (Dumoulin and Wandell, 2008). Thus, when we use stimulus location as our axes, the activity profiles will cluster around the axes, and their distribution will be distinctly non-Gaussian. This does not mean that assumption of Gaussian distributed activity profiles (as made in PCM) would automatically lead to invalid inference. Independent of the shape of the distribution, the second moment of the activity profiles determines which stimulus features can be read out from the neuronal code. Only when the read out from the area is restricted to be local, do higher order moments play a key role. The question of whether we can find evidence for clear non-Gaussian distributions of activity profiles, and what these may imply for the neural representations in these regions, is in highly interesting topic that warrants further study (Norman-Haignere et al., 2013).

## Footnotes

1 It is important to keep in mind that the second moment matrix (and hence correlations or distances) calculated from the pattern estimates **Û** do not equal those derived directly from our estimate **G**(*θ*). This is because the former do not reflect the uncertainty in the activation estimates, which plays a role in determining the second moment. In short, the second moment of the mean estimates does not equal the mean second moment estimate. Therefore, the activity estimates should only be used to visualize the spatial arrangement of activity patterns, not to visualize their representational structure.

